# Time-dependent synergy for multi-agent anticancer therapy

**DOI:** 10.1101/2020.10.08.332031

**Authors:** Krzysztof Fujarewicz, Krzysztof Łakomiec

## Abstract

A synergy between two therapeutic agents occurs when the effect of using both is greater than the sum of the effects of using each of them separately. There are several methods of detecting synergies in the literature, but they do not take into account the relationship between the times in which the factors act. In the article, we propose and justify the use of second-order sensitivity analysis as a potential tool for the detection and visualization of this type of synergy. We test and illustrate the proposed approach using four exemplary models of combined radio-chemotherapy of cancer.

## 1 Introduction

In modern medicine, combination therapy is very often used, in which two or more therapeutic agents are used simultaneously. A very desirable effect then is the so-called synergistic effect, in which the therapeutic effect is not only the sum of the effects of each of them separately, but is additionally enhanced.

Several general methods have been established to evaluate therapeutics agents combinations. The most commonly used are based on so-called isobolograms [1]. The isobologram shows relationship between drug dose and a biological response. Usually the isobologram generates a curve which can be used to distinguish simple additivity from synergy (or antagonism) effect. A little modification to the isobologram method is the Tumor Static Concentration curve where involves plasma concentration as opposed to the dose [2]. Additionally, in recent years several machine learning approach has been used for identifying synergistic drug combinations [3, 4].

However methods presented above does not provide any information about the effect of changing of the synergy over time. To fill this gap we present a method to study the synergy effect using local sensitivity analysis. Sensitivity analysis can be viewed as study of how the change of system’s input affects the change of particular system’s output. This kind of analysis has been used in our previous works to study the behaviour of complex biomedical models [5], parameter estimation process of mathematical models described by various type of differential equations [6–9], or during control optimization tasks [10, 11].

In this work we propose a numerical method to check the existence of the synergistic effect in models of combined therapy described by ordinary differential equations (ODE). Proposed method is based on the concept of second-order sensitivity analysis, and assumes numerical calculation of objective function’s second derivative. Value of this derivative informs us about the existence of the synergistic effect in the analysed model, and it depends on the form of the objective function used in analysis. If the objective function reflects the therapeutic effect (i.e. both raises together) then positive value of second order derivatives of the objective function indicates the existence of the synergistic effect.

Let us assume that the effect of two factors *x*_1_ and *x*_2_ is expressed by given function *f*(*x*_1_, *x*_2_). Then we say that there exists synergistic effect (synergy) between *x*_1_ and *x*_2_ when

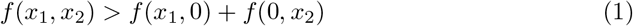

Now, let us treat as an effect the *change* of the function Δ*f* caused by changes Δ*x*_1_ and Δ*x*_2_. Then the synergy between Δ*x*_1_ and Δ*x*_2_ exists when

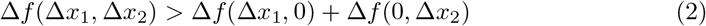

or alternatively when

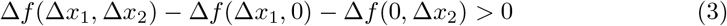

In this article we will describe the method of detection of the second *incremental* type of synergy denoted by (2) and (3).

## 2 Second order sensitivity analysis for synergy detection

The behavior of the model described by the system of ordinary differential equations depends on the values of constant parameters (including the initial values) and on the signals (variable parameters) affecting the system. Taking this into account, three different second order sensitivity analyzes are possible — a sensitivity analysis taking into account changes in:

- parameter and parameter, i.e. classical 2nd order parametric sensitivity analysis,
- parameter and signal,
- signal and signal.

Two latter cases above are *time-dependent sensitivities*. In the subsections below, we present how to derive the sensitivities of all three types. At the same time, we describe how the obtained results should be interpreted in the context of the synergistic effect.

### 2.1 Parametric second order sensitivity

Let us assume that a given objective function *J* depends on solution (trajectories) of the model having two parameters *p*_1_ and *p*_2_. The coefficient 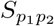 will stand for the second derivative of the objective function *J* with respect to the parameters *p*_1_ and *p*_2_:

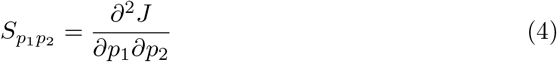

To approximate the coefficient 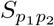 one can use the finite difference method:

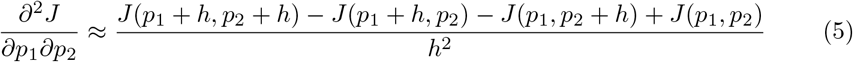

where *p*_1_, *p*_2_ are nominal values of analyzed parameters, *h* is a step of the finite difference approximation.

#### 2.1.1 The value of the second order sensitivity and its connection with synergy

The coefficient 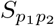 is a *linear operator* and may be used to calculate how the “small” change of one parameter influence the first order sensitivity with respect to the second parameter:

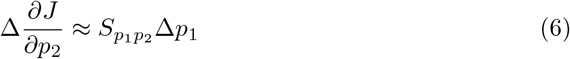

Hence for positive value of 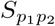 and positive Δ*p*_1_ the sensitivity of *J* with respect to *p*_2_ increases.

Using other words we can say that when 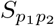 is positive then change of one parameter *sensitizes* the objective function *J* to another parameter. The relation is symmetric and we have also:

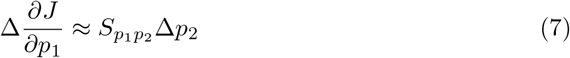

The relationship of second order sensitivity with synergy can be also demonstrated on another way. Let us rewrite the numerator of the right size of (5) as follows

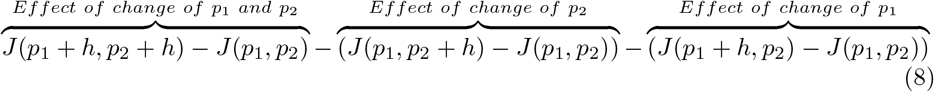

When we compare (8) with the synergy condition (3) then it becomes clear that the positive sign of *S*_*pp*_ indicates the existence of synergy.

### 2.2 Time-dependent second order sensitivity

As mentioned above, two types of time-dependent second order sensitivity can be distinguished:

- sensitivity with respect to parameter and signal,
- sensitivity with respect to signal and signal.

#### 2.2.1 Sensitivity with respect to parameter and signal

Let us assume that the objective function *J* is influenced by a time-varying signal *u*(*t*) and a constant parameter *p*. Second derivative ^1^ of the objective function *J* with respect to the input signal *u*(*t*) and parameter *p* is a linear operator represented as a following function of time:

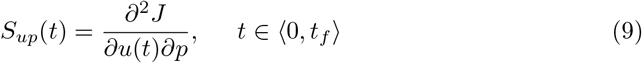

which can be used to calculate change of the sensitivity of *J* with respect to the parameter *p* as a scalar product of *S*_*up*_ and change of *u*(*t*):

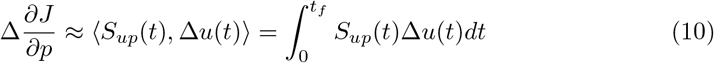

or, on the other hand, to calculate the change of the sensitivity of *J* with respect to the signal *u*(*t*):

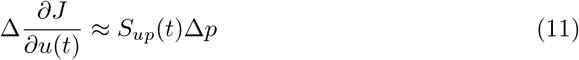

To approximate the function *S*_*up*_ at a fixed time *t*\* one can use the finite difference method:

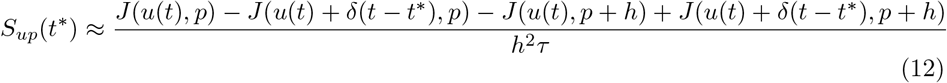

where *δ*(*t*) is a rectangular pulse function which is equal to *h* for 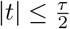 and is equal to 0 otherwise. *u*(*t*) and *p* are nominal input signal and parameter, *h* is a step of the finite difference approximation.

An example of second order sensitivity function *S*_*up*_(*t*) with respect to one signal and one parameter is shown on Fig. 1. On this plot can be seen that the synergy effect depends on time — at time *t* equal to 3 the sensitivity function is negative, in other case, at time equal to 7, the sensitivity is greater than zero. That means if we increase the value of signal *u* at time 3 we will decrease the sensitivity of the objective function *J* to the parameter *p*. In the second case when we increase the value of signal *u* at time 7 we will increase the sensitivity of objective function *J* with respect to parameter *p*.

**Fig 1.**
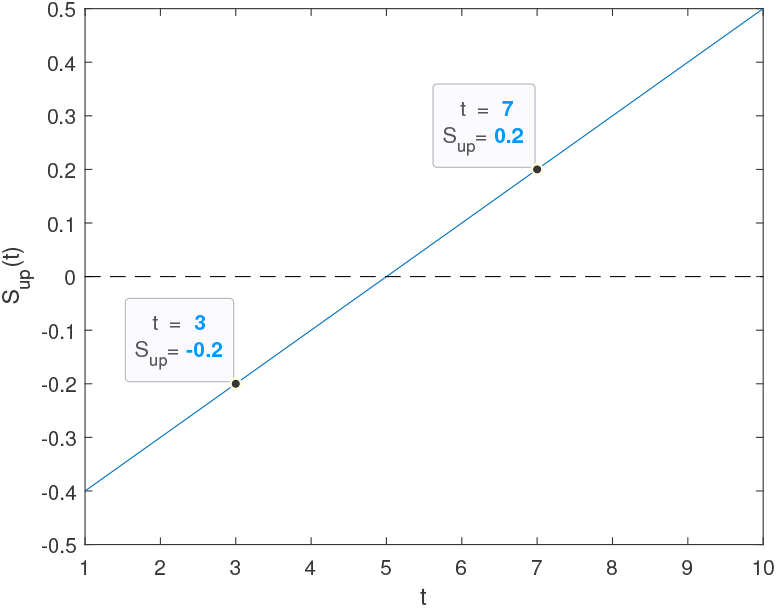
Example of second order sensitivity function with respect to one parameter *p* and one signal *u*(*t*).

On the other hand when we increase the parameter *p* then we decrease the sensitivity of *J* with respect to *u*(*t*) for *t <* 5 and increase this sensitivity for *t >* 5.

#### 2.2.2 Sensitivity with respect to two signals

Let us assume that the objective function *J* is influenced by two time-varying signals *u*_1_(*t*) and *u*_2_(*t*). Second order (Fréchet) derivative of the objective function *J* with respect to the input signals *u*_1_(*t*) and *u*_2_(*t*) can be represented as a following function of two time variables *t*_1_, *t*_2_:

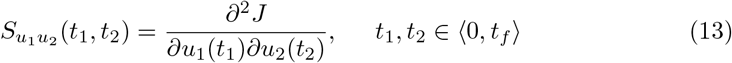

As previously defined two types of sensitivity 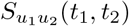 is a linear operator and may be used to obtain change of the sensitivity of *J* with respect to one signal caused by change of another signal. The value of 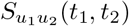 for fixed times *t*_1_ = *t*\* and *t*_2_ = *t*\*\* inform us how the change of *u*_1_(*t*\*) *sensitizes J* to changes of *u*_2_(*t*\*\*) and *vice versa*.

Let 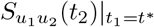 be an intersection of (13) for fixed *t*_1_ = *t*\* and 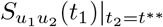 be an intersection of (13) for fixed *t*_2_ = *t*\*\*. Both intersections are functions of single time variable and can be used to calculate the change of the sensitivity of *J* with respect to *u*_1_(*t*\*) caused by change of whole signal *u*_2_(*t*_2_):

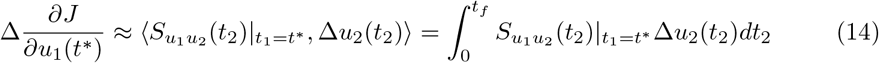

or to calculate the change of the sensitivity of *J* with respect to *u*_2_(*t*\*\*) caused by change of whole signal *u*_1_(*t*_1_):

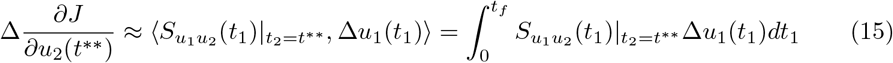

To approximate the function 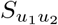 one can use the finite difference method:

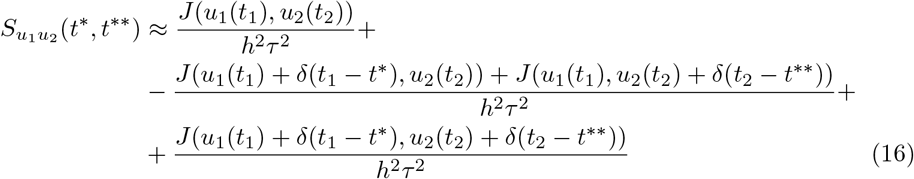

where *u*_1_(*t*_1_), *u*_2_(*t*_2_) are nominal input signals and *δ*(*t*) is, as previously, the rectangular step function of amplitude *h* which lasts for 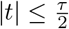.

An example of second order sensitivity function 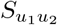 with respect to two signals signals is shown on center panel in Fig. 2. It is clearly visible that this sensitivity function depends on times *t*_1_ and *t*_2_. It can be interpreted as follows: if we increase the signal *u*_1_ at time *t*\* = 20 then the sensitivity of objective function *J* with respect to the signal *u*2 will change as the cross-section shown on the left panel of Fig. 2. In other case when we increase the signal *u*_2_ at time *t*\*\* = 80 then the sensitivity of the objective function *J* with respect to the signal *u*_2_ will change as the cross-section shown on the right panel of Fig. 2. In other case when we increase the signal *u*_2_ at time *t*\*\* = 80 then the sensitivity of the objective function *J* with respect to the signal *u*_1_ will change as the cross-section shown on the right panel of Fig. 2.

**Fig 2.**
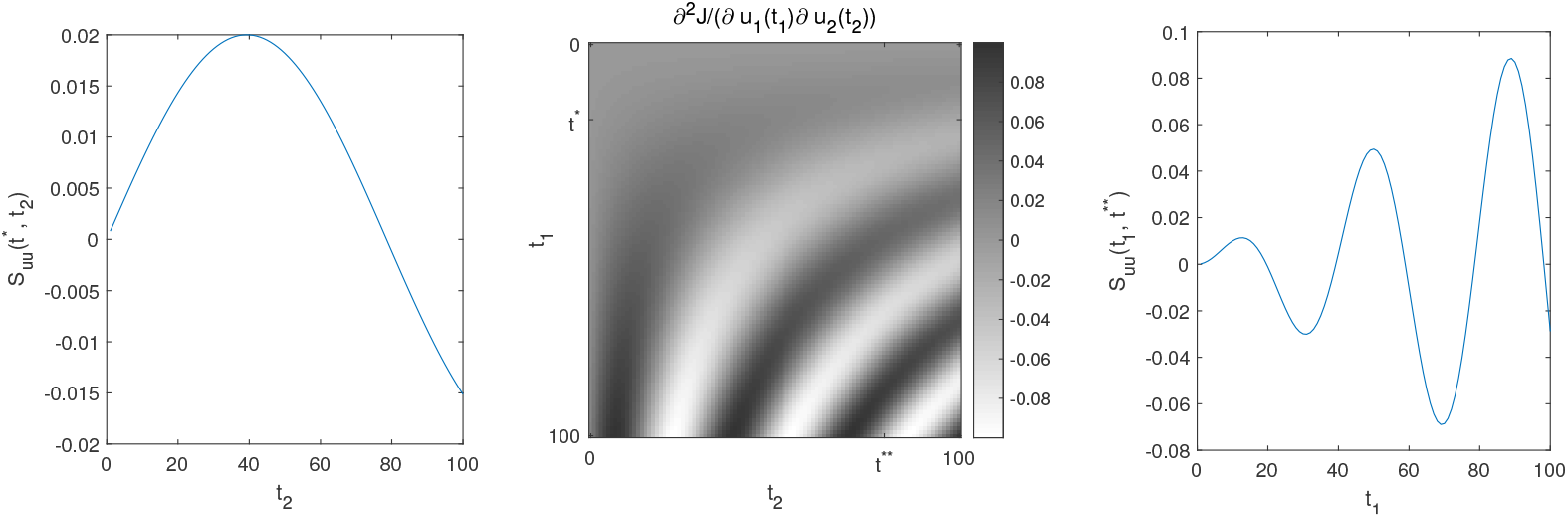
*Center panel:* Example of second order sensitivity of the objective function *J* with respect to two signals *u*_1_(*t*_1_), *u*_2_(*t*_2_). *Left panel:* Cross-section of the example sensitivity function at time *t*_1_ = *t*\* = 20, *Right panel:* Cross-section of the example sensitivity function at time *t*_2_ = *t*\*\* = 80.

## 3 Analysed models of radio-chemotherapy

We used presented method to study of the synergy effect in the following models of radio-chemotherapy of a tumor:

- Model 1 First model was acquired from work [12] and is defined as following ordinary differential equation:

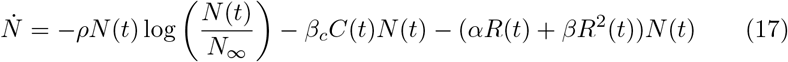

where *N* is the tumor size, *N*_∞_ is the carrying capacity of the tumor, *ρ* is the proliferation coefficient, *β*_*c*_ is a coefficient of sensitivity tumor to chemotherapy, and *α, β* are parameters of the linear-quadratic (LQ) response model of tumor to radiotherapy. The input signals *R*(*t*) and *C*(*t*) represent the dose of radiotherapy and chemotherapy at time *t*.
- Model 2 Second model was acquired from work [13] and is also defined as one ordinary differential equation:

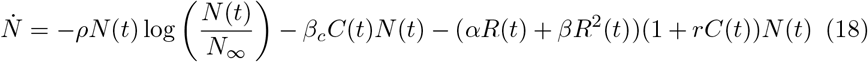 Model introduces the synergy between radio and chemotherapy, and coefficient *r* indicating the strength of this effect.
- Model 3 Third analyzed model is similar to the second but introduces additionally the effect of pharmacokinetics (*γ*(*t*)). Therefore the third model is described as a system of two ordinary differential equations:

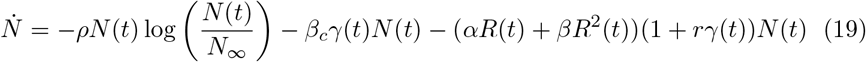

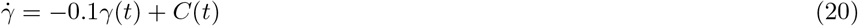
- Model 4 The last analyzed model in this work introduces the mechanism of DNA double strand breaks repair in the linear-quadratic model according to the work [14]. Therefore it is described as a system of three ordinary differential equations:

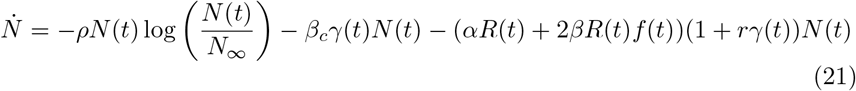

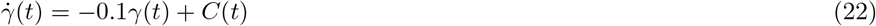

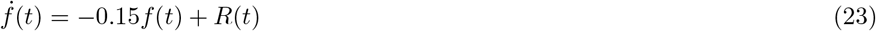

### 3.1 Nominal parameters, models solver and simulations

Nominal parameters of the analyzed models were acquired from work [13]: *ρ* = 7e-5, *N*_∞_ = 1.735e4, *β*_*c*_ = 1.4e-2, *α* = 3.98e-2, *β* = 3.98e-3, *r* = 0.1. Initial conditions for the models were set as *N* (0) = 3e3, *γ*(0) = *f*(0) = 0. Nominal input signals *R*(*t*), *C*(*t*) are assumed to be uniformly distributed with value equal to 0.005 at any time *t*: *R*(*t*) = 0.005, *C*(*t*) = 0.005.

To solve the models we used function *ode15s* from MATLAB environment. To avoid numerical errors during analysis we set the maximum step size for the solver to 1e-3. The final time *t*_*f*_ of simulations during analysis was set to 20. Simulation results of the models with nominal values of parameters and inputs are shown in Fig. 3.

**Fig 3.**
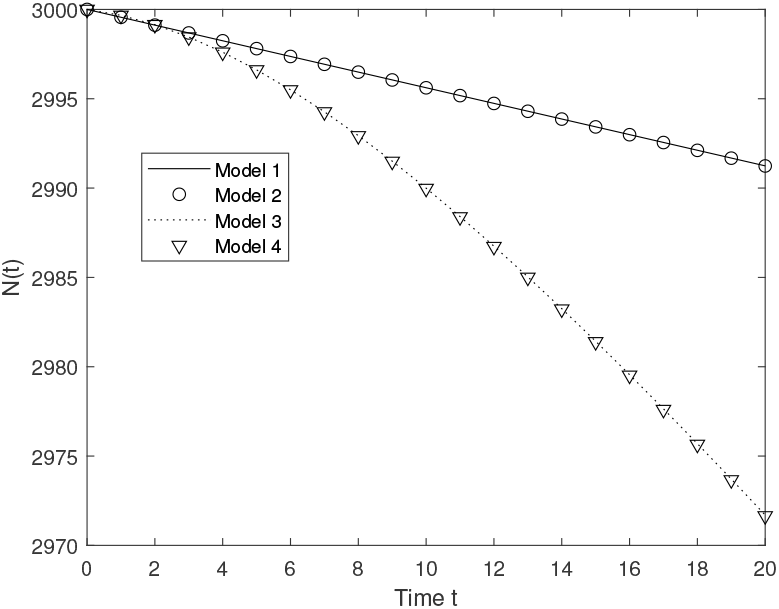
Simulation of the models with nominal values of parameters and signals.

## 4 Results

All presented results of the second order sensitivity analysis are obtained using Finite Difference Method (FDM) with the step sizes *h* and *τ* set to 1e-3 and 0.2 respectively. Due to numerical nature of described method a threshold value must be defined below which the resulting value are treated as zero. In our analysis we set the threshold value to 5e-6.

### 4.1 Objective function

Objective function for synergy analysis in the models was defined as:

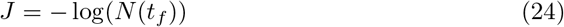

where *N*(*t*_*f*_) is the tumor size at final time of simulation. The sign “−” is related to the requirement that larger value *J* should correspond to stronger therapeutic effect. In other words, therapy optimization means maximizing the objective function.

### 4.2 Parametric second order sensitivity

Results of the parametric second order sensitivity analysis using equation (1) for Models 1–4 are shown in tables 1, 2, 3, 4 respectively. The value in a table row marked by *∂p*_1_ and a column marked by *∂p*_2_ stands for 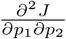.

**Table 1.** Results of the parametric second order sensitivity analysis for model 1.

| | $\partial\rho$ | $\partial N_\infty$ | $\partial\beta_c$ | $\partial\alpha$ | $\partial\beta$ |
| --- | --- | --- | --- | --- | --- |
| $\partial\rho$ | 757.5789 | -0.0012 | -1.0937 | -1.0937 | -0.0055 |
| $\partial N_\infty$ | -0.0012 | 0 | 0 | 0 | 0 |
| $\partial\beta_c$ | -1.0937 | 0 | 0 | 0 | 0 |
| $\partial\alpha$ | -1.0937 | 0 | 0 | 0 | 0 |
| $\partial\beta$ | -0.0055 | 0 | 0 | 0 | 0 |

**Table 2.**
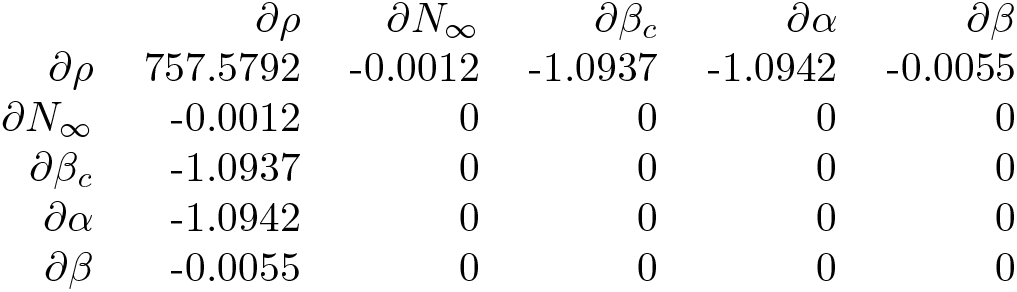
Results of the parametric second order sensitivity analysis for model 2.

**Table 3.**
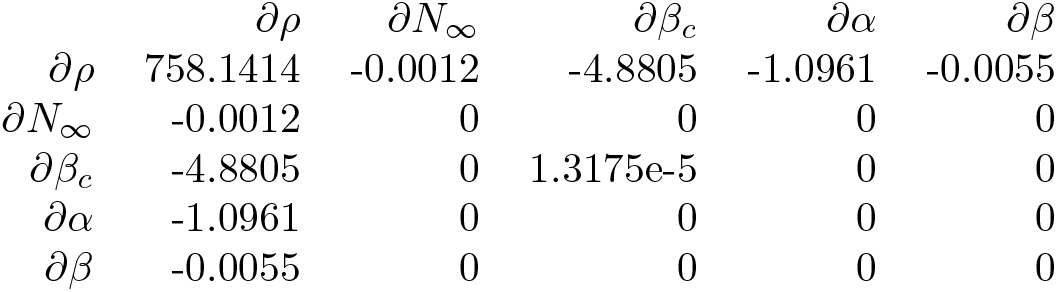
Results of the parametric second order sensitivity analysis for model 3.

| | $\partial\rho$ | $\partial N_\infty$ | $\partial\beta_c$ | $\partial\alpha$ | $\partial\beta$ |
| --- | --- | --- | --- | --- | --- |
| $\partial\rho$ | 758.1414 | -0.0012 | -4.8805 | -1.0961 | -0.0055 |
| $\partial N_\infty$ | -0.0012 | 0 | 0 | 0 | 0 |
| $\partial\beta_c$ | -4.8805 | 0 | 1.3175e-5 | 0 | 0 |
| $\partial\alpha$ | -1.0961 | 0 | 0 | 0 | 0 |
| $\partial\beta$ | -0.0055 | 0 | 0 | 0 | 0 |

**Table 4.** Results of the parametric second order sensitivity analysis for model 4.

| | $\partial\rho$ | $\partial N_\infty$ | $\partial\beta_c$ | $\partial\alpha$ | $\partial\beta$ |
| --- | --- | --- | --- | --- | --- |
| $\partial\rho$ | 758.1430 | -0.0012 | -4.8805 | -1.0961 | -0.0409 |
| $\partial N_\infty$ | -0.0012 | 0 | 0 | 0 | 0 |
| $\partial\beta_c$ | -4.8805 | 0 | 1.3190e-5 | 0 | 0 |
| $\partial\alpha$ | -1.0961 | 0 | 0 | 0 | 0 |
| $\partial\beta$ | -0.0409 | 0 | 0 | 0 | 0 |

The highest values in all models were achieved by the derivative with respect to *∂ρ*^2^. Another interesting observation for model 1 is the values of derivatives with respect to *∂ρ∂β*_*c*_ and *∂ρ∂α* are equal. This could mean that there are similar mechanisms in the model for chemo and radiotherapy (nominal inputs signals *R*(*t*) and *C*(*t*) are equals, and there is a linear term describing effect of chemotherapy and a linear term in the LQ model).

### 4.3 Time-dependent second order sensitivity

#### Sensitivity with respect to parameter and signal

Results of the time-dependent second order sensitivity using equation (12) are shown on fig. 4, 5, 6. Results for models 1 and 2 are the same and are therefore presented in one common fig. 4. First row of panels in the figure presents the second order sensitivity of *J* with respect to individual parameters and the radiotherapy *R*(*t*). Second row of panels presents the same but for the chemotherapy C(t).

**Fig 4.**
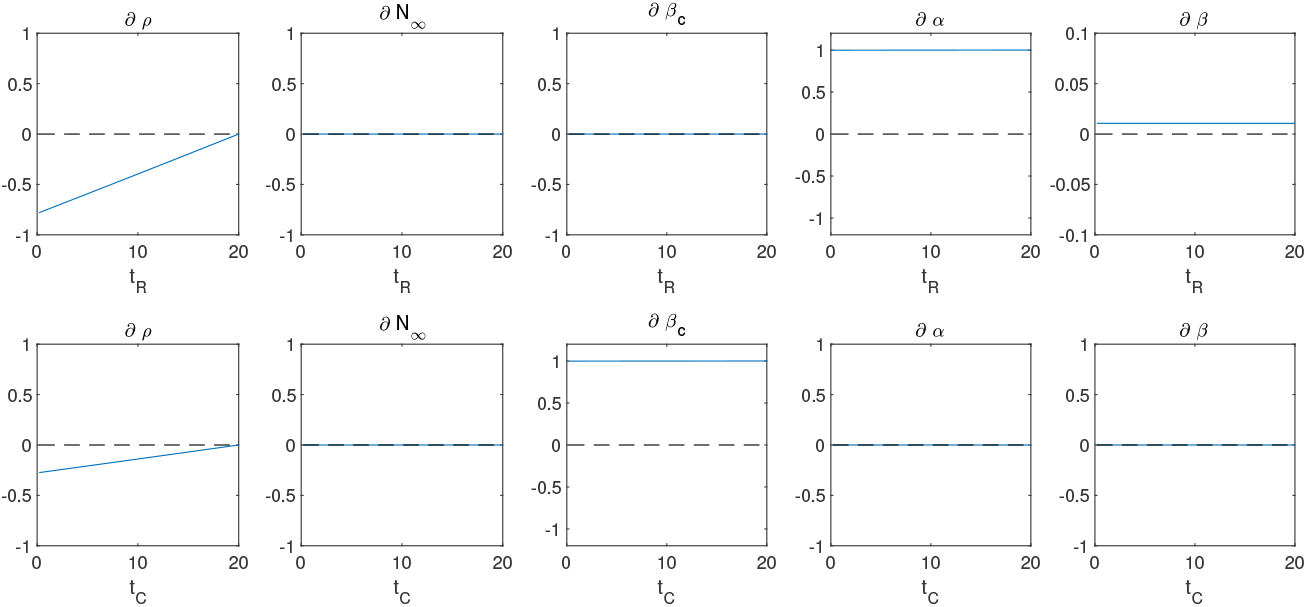
Results of the time-dependent second order sensitivity using equation 12 for models 1 and 2.

**Fig 5.**
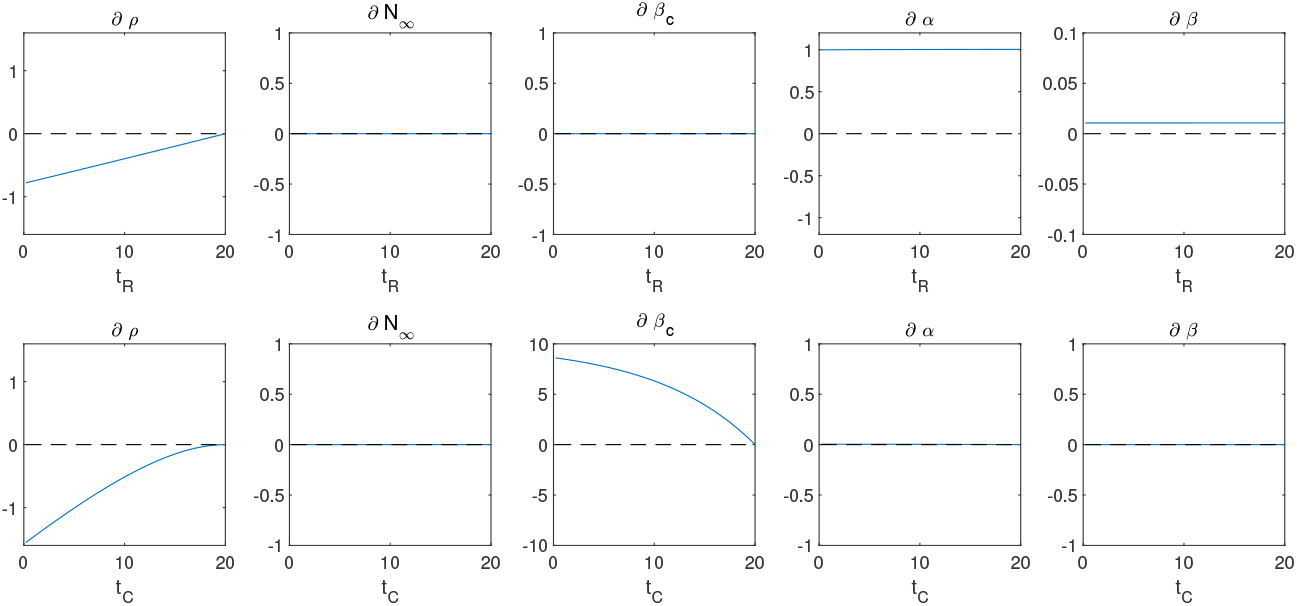
Results of the time-dependent second order sensitivity using equation 12 for model 3.

**Fig 6.**
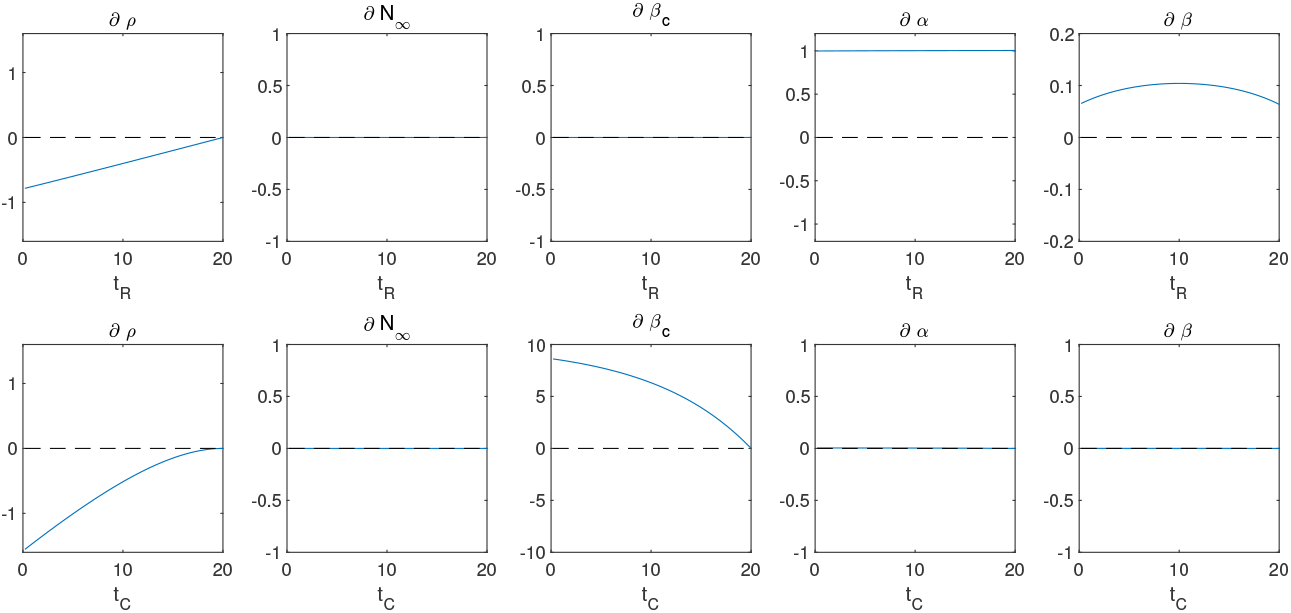
Results of the time-dependent second order sensitivity using equation 12 for model 4.

In all models there exists s a synergy between chemotherapy signal *C*(*t*) and parameter *β*_*c*_. Similarly, in all analysed models there is a synergy between radiotherapy signal *R*(*t*) and parameters *α*, *β*. In models 3 and 4 these synergies changing in time due to additional equations describing pharmacokinetics and DNA repair.

#### Sensitivity with respect to two signals

Results of the time-dependent second order sensitivity using equation (16) are shown on figs. 7, 8. The derivatives with respect to *C*(*t*)^2^ are equal to zero in all analyzed models, indicating that there is no synergy in the action of chemotherapy alone.

**Fig 7.**
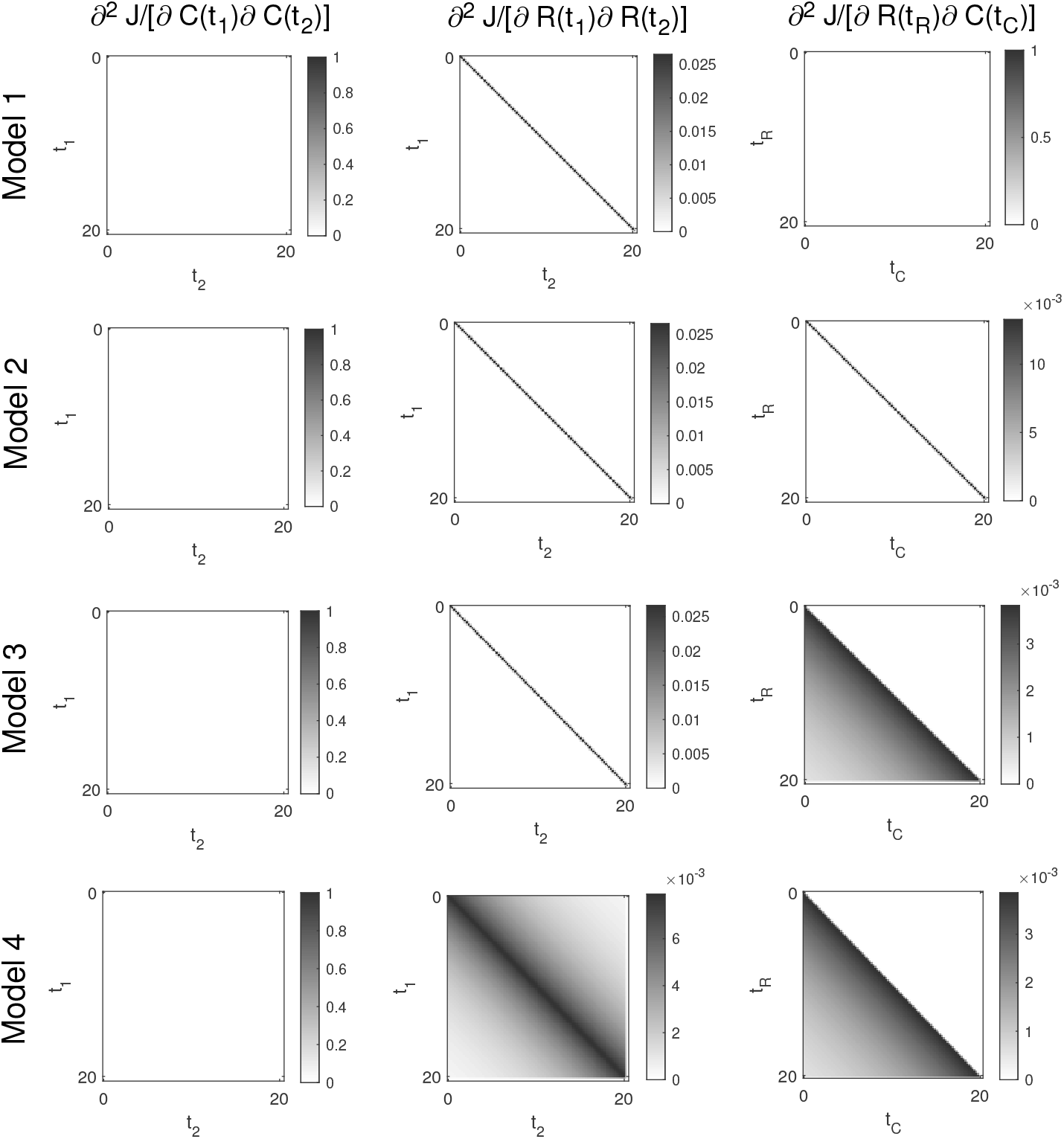
Results of the time-dependent second order sensitivity using equation (16).

**Fig 8.**
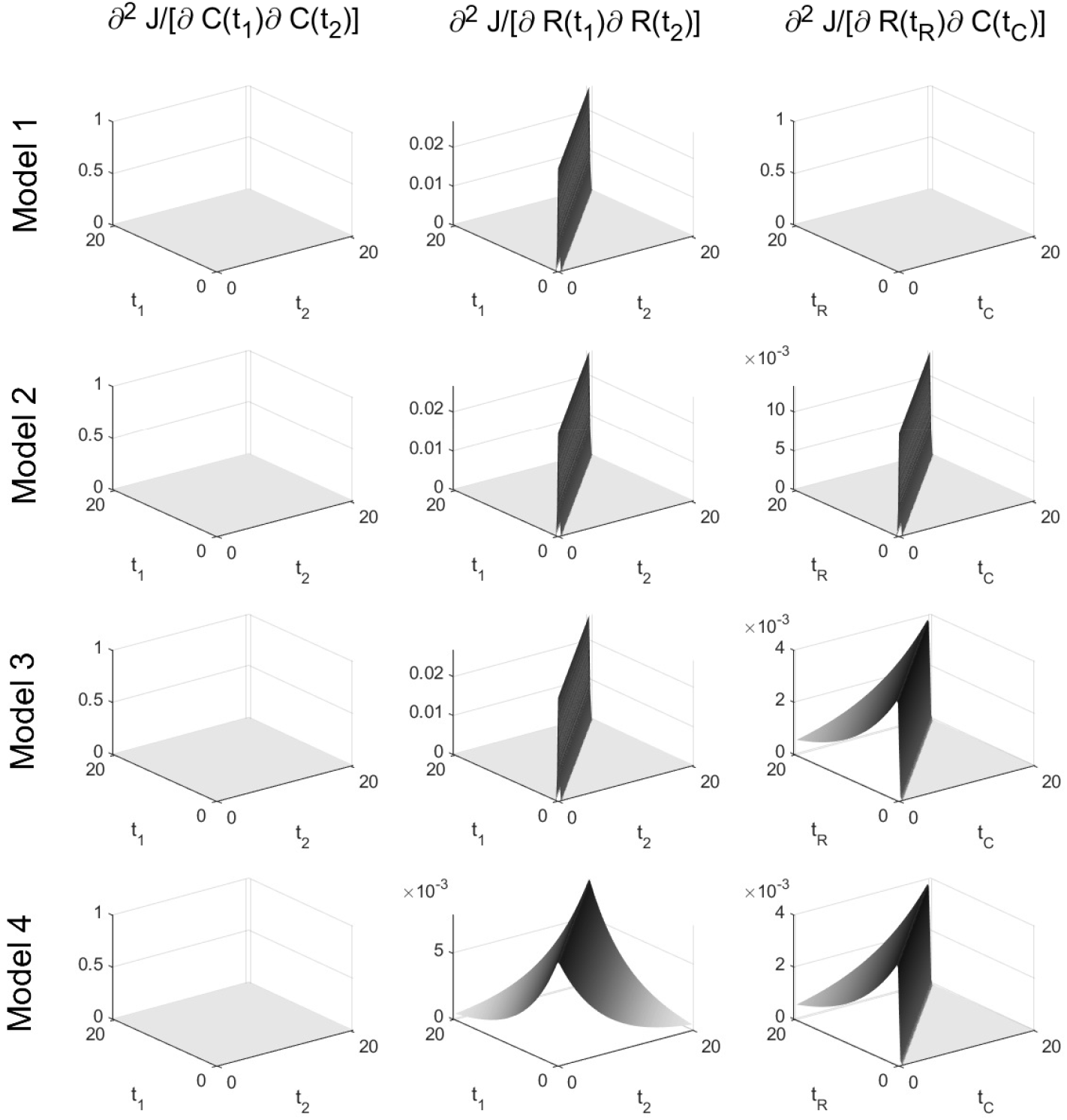
Results of the time-dependent second order sensitivity using equation (16).

For radiotherapy alone there is a synergy effect in all four models. This is due to the quadratic term in LQ model. For models 1-3 the synergy for radiotherapy alone is only visible only when we change at one time *t* the signal *R*(*t*), but for model 4 due to DNA repair mechanism this effect is spreading over time.

The synergy effect between radio and chemotherapy is visible only in models 2-4. In model 1 the derivative with respect to *R*(*t*)*C*(*t*) is zero because there is no synergy term between these two therapies. In model 2 the synergy between radio and chemotherapy only occurs when we changing the signals *R*(*t*) and *C*(*t*) at the same time. Models 3 and 4 due to the pharmacokinetics effect only shows the synergy effect between these two treatments when chemotherapy is added before radiotherapy.

## 5 Discussion

Results obtained for analyzed models of radio-chemotherapy shows that synergy effect can be manifested in many ways, and classical isobologram based methods for study this effect may be insufficient. It appears that the important thing in synergy between two therapeutics agents may be study how this effect can change in time during therapy. Deepening the knowledge about this aspect of synergy may results in better plans for therapies against tumors.

## 6 Conclusion

Presented in this work method can be used to study how the synergy effect changing in time in mathematical models describing two therapeutics agents. However method can be used to any mathematical models described by ordinary differential equations. It is based on second-order sensitivity analysis and assume numerical approximation of objective function’s second order derivatives. The main advantages of this method is simplicity, using only finite difference approximation. Due to numerical nature user must only set the valid threshold value and the step size.

## Acknowledgments

This work was supported by the Polish National Science Centre under grant DEC-2016/21/B/ST7/02241 (K.F.), and by the Silesian University of Technology under grant 02/040/BKM20/0007 (K.Ł.).

Calculations were performed using the infrastructure supported by the computer cluster Ziemowit (www.ziemowit.hpc.polsl.pl) funded by the Silesian BIO-FARMA project No. POIG.02.01.00-00-166/08 and expanded in the POIG.02.03.01-00-040/13 in the Computational Biology and Bioinformatics Laboratory of the Biotechnology Centre at the Silesian University of Technology.

## Footnotes

1 More mathematically precisely: second order Fréchet derivative

## References

1. Huang R-y, Pei L, Liu Q, Chen S, Dou H, Shu G, Yuan Z-x, Lin J, Peng G, Zhang W and Fu H (2019) Isobologram Analysis: A Comprehensive Review of Methodology and Current Research. Front. Pharmacol. 10:1222. doi: 10.3389/fphar.2019.01222

2. Cardilin T, Almquist J, Jirstrand M, Zimmermann A, El Bawab S, Gabrielsson J. Model-Based Evaluation of Radiation and Radiosensitizing Agents in Oncology. CPT Pharmacometrics Syst Pharmacol. 2018;7(1):51–58. doi:10.1002/psp4.12268

3. Gayvert KM, Aly O, Platt J, Bosenberg MW, Stern DF, Elemento O (2017) A Computational Approach for Identifying Synergistic Drug Combinations. PLoS Comput Biol 13(1): e1005308

4. Julkunen, H., Cichonska, A., Gautam, P. et al. Leveraging multi-way interactions for systematic prediction of pre-clinical drug combination effects. Nat Commun 11, 6136 (2020)

5. Łakomiec K., Kurasz K., Fujarewicz K. (2019) Sensitivity Analysis of Biomedical Models Using Green’s Function. In: Pietka E., Badura P., Kawa J., Wieclawek W. (eds) Information Technology in Biomedicine. ITIB 2018. Advances in Intelligent Systems and Computing, vol 762. Springer, Cham.

6. Łakomiec, K., Kumala, S., Hancock, R., Rzeszowska-Wolny, J., Fujarewicz, K.: Modeling the repair of DNA strand breaks caused by *γ*-radiation in a minichromosome. Phys. Biol. 11(4), 045003 (2014).

7. Łakomiec, K., Fujarewicz, K.: Parameter estimation of non-linear models using adjoint sensitivity analysis. In: Advanced Approaches to Intelligent Information and Database Systems, Studies in Computational Intelligence, vol. 551, pp. 59–68.

8. Jakubczak, M., Fujarewicz, K.: Application of adjoint sensitivity analysis to parameter estimation of age-structured model of cell cycle. In: Pietka, E., Badura, P.,Kawa, J., Wieclawek, W. (eds.) Advances in Intelligent Systems and Computing, vol. 472, pp. 123–131. Springer (2016)

9. Fujarewicz, K., Łakomiec, K.: Parameter estimation of systems with delays via structural sensitivity analysis. Discr. Continuous Dyn. Syst. Ser. B 19(8), 2521–2533 (2014).

10. Fujarewicz, K., Łakomiec, K.: Adjoint sensitivity analysis of a tumor growth model and its application to spatiotemporal radiotherapy optimization. Math. Biosci. Eng. 13(6), 1131–1142 (2016).

11. Fujarewicz K, Łakomiec K: Spatiotemporal sensitivity of systems modeled by cellular automata. Mathematical Methods in the Applied Sciences, 2018, 41(18), p. 8897–8905

12. P. Bajger, K. Fujarewicz, and A. Swierniak. Effects of pharmacokinetics and DNA repair on the structure of optimal controls in a simple model of radio-chemotherapy. 2018 23rd International Conference on Methods and Models in Automation Robotics (MMAR), 2018. doi:10.1109/MMAR.2018.8485901.

13. P. Bajger, Krzysztof Fujarewicz, Andrzej Świerniak., Optimal control in a model of chemotherapy-induced radiosensilisation.Math. Appl. 2019 vol. 47 iss. 1, p. 81–91

14. R.K. Sachs, L.R. Hlatky, P. Hahnfeldt, Simple ODE models of tumor growth and anti-angiogenic or radiation treatment, Mathematical and Computer Modelling, Volume 33, Issues 12–13,2001, Pages 1297–1305,

